# Physically stressed bees expect less reward in an active choice judgement bias test

**DOI:** 10.1101/2023.10.06.561175

**Authors:** Olga Procenko, Jenny Read, Vivek Nityananda

## Abstract

Emotion-like states in animals are commonly assessed using judgment bias tests, which measure responses to ambiguous information. A few studies have recently used these tests to argue for the presence of emotion-like states in insects. However, the results from most of these studies could have other explanations, including changes in motivation and attention. To control for these explanations, we therefore developed a novel judgment bias test, requiring bumblebees to make an active choice indicating their interpretation of ambiguous stimuli. Bumblebees were trained to associate high or low rewards, delivered in two different reward chambers, with distinct colors. Two groups of bees were then physically stressed by shaking or trapping, while the third group served as a control. We subsequently presented the bees with ambiguous colors between the two learnt colors and noted which reward chamber they chose. When presented with ambiguous colors, stressed bees were less likely than control bees to enter the reward chamber previously associated with high reward. We modelled bee behavior using signal detection and drift diffusion models and showed that control bees and stressed bees were, respectively, more likely to respond optimistically and pessimistically to ambiguous cues. The signal detection model further showed that the behavior of stressed bees was explained by a reduction in their prior expectation of high rewards. Our findings thus provide strong evidence for emotion-like states in bees and suggest that their stress-induced pessimistic behavior is explained by a reduced expectation of higher rewards.

## Introduction

The presence of emotions in non-human animals is much debated and can have important societal implications. While most research on animal emotions has focused on vertebrates (1,2), a handful of recent studies have explored analogous states in insects (3–8). In these studies, emotions are defined as valenced brain states triggered by both internal and external stimuli and comprising subjective, behavioral, physiological and cognitive components. Research on emotion-like states in insects has primarily relied on judgement bias tests, a method initially developed for assessing affective states in rats (9). These tests assess how animals respond to ambiguous stimuli. An animal typically is trained to associate one stimulus with a reward and another with a lack of reward or punishment. It is then tested with an ambiguous stimulus that is in-between the two learnt stimuli. Animals that respond as if this stimulus indicates a reward are considered optimistic, while those that respond as if the stimulus indicates a lack of reward or punishment are considered pessimistic.

Judgement bias tests have been used in five studies on insects, including on honeybees, bumblebees and fruit flies (3–7). Some of these studies showed that physical agitation can reduce the response of bees and flies to ambiguous odors (3–5). Others showed that bees are quicker to fly towards (6) and more likely to choose (7) ambiguous visual stimuli after encountering an unexpected reward of sucrose solution, suggesting optimistic behavior. While these results parallel results from studies of emotions in vertebrates, other explanations have also been suggested, including changes in motivation or the ability to learn training cues (10,11).

One factor that complicates the interpretation of these results is that the majority of insect studies so far have utilized a go/no-go type of judgment bias task. Here, the animal is trained to respond to a positive stimulus (“go”) and suppresses the response to a negative one (“no-go”). When faced with an ambiguous stimulus, responding (“go”) or suppressing (“no-go”) a response is thought to reflect optimistic and pessimistic judgements, respectively. While this approach has been successfully used in many studies across taxa (12–14), there are concerns associated with this paradigm. Firstly, the suppression of a response could result from a general reduction in activity and motivation rather than a judgment bias (12). A reduction of responses could also indicate merely an absence of response (omission) rather than a deliberate choice (17,18). Finally, an animal may fail to attend to or detect a stimulus, and in such cases, the lack of a response could be mistakenly attributed to a pessimistic judgment (14,19). Without a test that can address these issues, we currently do not have strong evidence of emotion-like states in insects. In addition, we lack models for the mechanisms underpinning the observed behaviors – though recent work has proposed that judgement biases in bees can arise from shifts in stimulus-response curves (7).

One way of reducing the likelihood of confounds is to use an active choice judgment bias test (16,17,20). In contrast to the go/no-go task, the active choice paradigm requires the animal to make an active choice between two alternative responses. Animals might, for example, learn to move to one location in response to one stimulus and to another location when they see another stimulus. Since the animal must make a choice as a response, this type of judgment bias test eliminates the possible confounding factors of the “go/no-go” paradigm, increasing validity and ease of interpretation.

We therefore used an active choice type of judgment bias test to rigorously assess judgement biases in bumblebees *(Bombus terrestris)*. Bees had to choose between two rewarding locations depending on the stimulus displayed, clearly signaling their judgement when faced with ambiguous stimuli by moving to one of the two locations. To induce negative affective states, we used two types of manipulations simulating predatory attacks - shaking and trapping by a robotic arm. These manipulations have previously been shown to be associated with cognitive and physiological changes (4,21,22). In addition, to further understand the mechanisms underlying our behavioral results, we applied drift diffusion and signal detection modelling frameworks to the data. We used these frameworks to test whether physical agitation affected the prior expectation of a reward in bees or their ability to distinguish between stimuli due to shifts in stimulus-response curves.

## Materials and Methods

### Animals and experimental set-up

All experiments were run on female worker bumblebees (*Bombus terrestris*) obtained from a commercial supplier (Koppert, UK). We transferred the bumblebees to one chamber of a bipartite plastic nest box (28.0 × 16.0 × 12.0 cm). We covered the other chamber of the nest box with cat litter to allow bees to discard refuse. The nest box was connected via a transparent acrylic tunnel (56.0 × 5.0 × 5.0 cm) to a flight arena (110.0 × 61.0 × 40.0 cm) with a UV-transparent Plexiglas® lid and lit by a lamp (HF-P 1 14-35 TL5 ballast, Philips, The Netherlands) fitted with daylight fluorescent tubes (Osram, Germany). When not part of an experiment, bees were fed with ∼ 3 g of commercial pollen daily (Koppert B. V., The Netherlands) and provided sucrose solution (20% w/w) ad libitum. Although invertebrates do not fall under the Animals (Scientific Procedures) Act, 1986 (ASPA), the experimental design and protocols were developed incorporating the 3Rs principles. Housing, maintenance, and experimental procedures were non-invasive and were kept as close as possible to the natural living conditions of the animals.

Visual stimuli were solid colors covering the entire display of an LED monitor (Dell U2412M, 24“, 1920 x 1200 px) and controlled by a custom-written MATLAB script (MathWorks Inc., Natick, MA, USA) using the PsychToolbox package (34). We measured the spectral reflectance of all colors used in the experiment using an Ocean Optics Flame reflectance spectrophotometer (Ocean Optics Inc., Florida, USA). The perceptual positions of the colors in the bee color hexagon space (Fig. 1B) were calculated using the spectral reflectance measurements and spectral sensitivity functions for *Bombus terrestris* photoreceptors (35,36).

**Figure 1.**
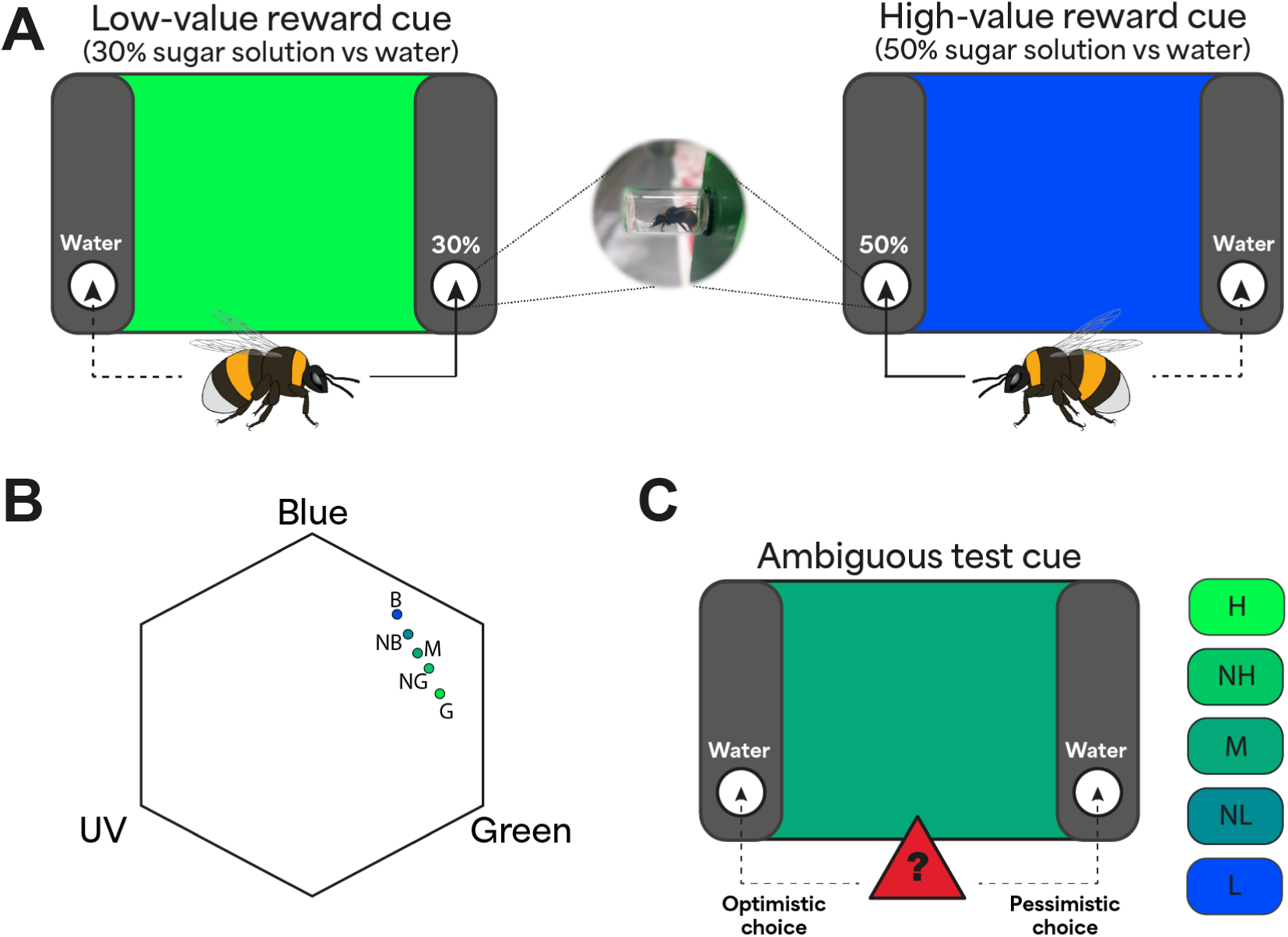
Experimental Protocol. **A)** Training phase. Bees were trained to associate two colors, green and blue, presented on an LED screen with different sugar rewards at different locations. The bees were presented one color at a time in pseudorandomized order. The figure depicts a training scenario with green associated with a low reward (30% sucrose solution) in the right chamber and blue with a high reward (50% sucrose solution) in the left chamber. The association between color, reward and location was counterbalanced across trials. Further details in the text. **B)** Cue colors plotted in bee color space (color cue: B, blue; NB, near blue; M, medium; NG, near green; G, green). The three vertices correspond to maximum excitation of photoreceptors sensitive to blue, green and ultraviolet (UV) light. The distance from the center to any vertex is 1 and the distance between points represents hue discriminability, with 0.1 being easily distinguishable. **C)** Judgement bias testing. The test phase consisted of five trials with different colors presented on the screen in a pseudorandom order (cue value: H, high; NH, near high; M, medium; NL, near low; G, low). The colors included the two conditioned colors and three ambiguous colors of intermediate value. In our example here, the screen shows the medium color with blue as the high-reward color (H) and green as the low-reward color (L), but this was counterbalanced across bees. Entering a chamber associated with a high reward during training was considered an optimistic choice, while entering a chamber associated with a low reward during training was deemed a pessimistic choice.

We positioned two vertical panels (40.0 × 8.0 cm) 8.5 cm in front of the right and left sides of the LED monitor, leaving the central area of the monitor open and visible. Each panel was equipped with an opening to place a reward chamber (7 ml glass vial, 10 mm inner diameter) 7 cm above the arena floor. Bees thus needed to fly from the arena entrance to the panels before entering the reward chamber. On each visit to the arena, the reward chambers were changed to ensure that pheromones and scent marks were not available during the next visit. In preparation for the next experimental day, all used chambers were washed in hot water and 70% ethanol and left to dry.

### Training procedure

Before the onset of training, bees were familiarized with both reward locations. A plastic cup was used to gently capture each bee. The opening of the cup was positioned so that it aligned with the entrance to the reward chamber, inside which the bee found a droplet of sucrose solution (0.2 ml, 30% w/w). We repeated the procedure equally on each side (left and right) without displaying any color on the LED screen. Individual bees that learnt the location of the reward and performed repeated foraging bouts were tagged for later identification using number tags (Thorne, UK). Tagging involved trapping each bee in a small marking cage, gently pressing it against the mesh with a sponge, and affixing the tag to the dorsal thorax with a small amount of superglue (Loctite Super Glue Power Gel).

In each training trial, we presented bees (n = 48) with one of two colors on the LED screen. The two colors used were green (RGB= 0, 255, 75) and blue (RGB= 0, 75, 225). When one of the colors was displayed, the bee was provided a high-value reward of 0.2 ml 50% (w/w) sucrose solution in one of the two chambers (e.g., on the left), and an equal amount of distilled water in the other chamber (e.g., on the right). In different trials, when the other color was displayed the bee was provided a low-value reward of 0.2 ml 30% (w/w) sucrose solution in the chamber opposite (e.g., on the right) to the one where, in the other trials, a high-reward was presented. Here again, an equal amount of distilled water would be present in the other chamber (e.g., on the left). Thus, on any given trial, the bee saw only one color and could encounter either the high or low reward (not both), with water on the unrewarding side. In addition, the locations of the high and low rewards were on opposite sides in their respective trials.

Across bees, the combinations of each color (green or blue), reward location (right or left) and reward type (high or low) were counterbalanced. Each bee encountered only one possible combination of each during training (e.g., green indicating a high reward on the left on half the trials, and blue indicating a low reward on the right on the other half). Trials presenting colors associated with high and low rewards were presented an equal number of times in a pseudorandom order, ensuring that no color was repeated more than twice in a row. To ensure that the bee entered the reward chamber fully to sample its content, we placed the droplets of solutions at the very end of the reward chamber (Fig. 1A). In all cases, the reward quantity of 0.2 ml allowed bees to fill their crop within a single reward chamber visit (37). We recorded a single choice on each trial, with a choice defined as a bee entering a chamber far enough to sample its content. Incidences of landing or partial entering (less than 1/3 of the body length) were not considered choices. Bees that reached the learning criterion (80% accuracy in the last 20 trials) continued to the test phase. 11 bees did not pass the initial conditioning test due to strong side biases. The last ten training trials were video recorded using a camera on a mobile phone (Huawei Nexus 6P phone 1440 × 2560 px, 120 fps) placed above the arena.

### Predatory attack simulation

We randomly assigned individual bees (n=48) that reached the learning criterion in the training phase to one of the three treatment groups. Two groups were subjected to manipulations which simulated predatory attacks and were predicted to change their affective state (4). One of these two treatments involved shaking the bee on a Vortex shaker (*Shaking*, n=16), while the other involved trapping the bee with a custom-made trapping device (*Trapping*, n=16). A third unmanipulated group served as a control (*Control*, n=16). The manipulations were applied to a bee before entering the arena for each test. Bees in the Control treatment were allowed to fly out into the flight arena without hindrance as in the training phase.

Each bee in the Shaking treatment was allowed to enter a custom-made cylindrical cage (40 mm diameter, 7.5 cm length). After entering, the bee was gently nudged down with a soft foam plunger until the distance between the plunger and the bottom of the cage was reduced to ∼3 cm. Once the plunger was secured, the cage with the bee was placed on a Vortex-T Genie 2 shaker (Scientific Industries, USA) and shaken at a frequency of 1200 rpm for 60 s. After shaking, the bee was released into the tunnel connecting the nest box and experimental arena via an opening on the top of the tunnel. The bee was released into the flight arena for testing as soon as it was ready to initiate a foraging bout.

Each bee in the Trapping treatment was trapped using a trapping device. This consisted of a soft sponge (3.5 × 3.5 × 3.5 cm) connected to a linear actuator system (rack and pinion). A micro-servo initiated the linear motion of the trapping device (Micro Servo 9g, DF9GMS), powered, and controlled by a microcontroller board (Arduino, Uno Rev 3). A custom-written script written in the Arduino Software (IDE) triggered an initial plunging movement of the trapping device, followed by release after three seconds. This permitted consistent trapping across all tested individuals. As in the Shaking treatment, the bee was released into the flight arena for testing as soon as it was ready to initiate a foraging bout.

### Judgement bias testing

The test phase consisted of five trials, each with a cue of a different color presented on the screen. The test colors were the two conditioned colors (green and blue), and three ambiguous colors of intermediate value between the two conditioned colors (near blue (RGB=0, 140, 150); medium (RGB= 0, 170, 120); near green (RGB= 0, 200, 100) (Fig. 1B). We classified the ambiguous colors as near-high, medium, and near-low cues depending on their distance to the high or low rewarding color for each bee. The color presentation order was pseudorandomized between all bees, so that the first test color was always one of the three ambiguous color cues. Within the test phase, all color cues (ambiguous and learnt) were not rewarded, i.e., both chambers contained 0.2 ml of distilled water. We classified the entry of a bee into a reward chamber as a choice. After it made the first choice, we gently captured the bee with a plastic cup and returned it to the tunnel connecting the nest and the arena. Between presentations of each of the five test cues, bees were provided refresher trials consisting of two presentations of each conditioned color with the appropriate reward at the correct location. All trials were video recorded for later video analysis using the camera of a mobile phone (Huawei Nexus 6P, 1440 × 2560 px, 120 fps). We obtained the latencies for the choices from the video analysis (see below).

### Measuring foraging motivation using ingestion rate

To assess if our manipulations changed feeding motivation in bees, we measured sugar reward ingestion rates. A separate group of bees (n=36) were pre-trained to forage from an elevated feeder consisting of the reward chamber used above with a 1.5 mL Eppendorf placed inside. After learning this location and completing five consecutive foraging bouts, bees were randomly allocated to one of three treatment groups as in the above experiment for the ingestion test (Control: n=12, Shaking: n=12, Trapping: n=12). The test consisted of a single foraging bout on a feeder with sucrose solution (∼1 ml, 50% w/w). The feeder was weighed before and immediately after the test bout to determine the mass of ingested solution using a Kern Weighing Scale ADB100-4 (Resolution: mg±0.001, Kern & Sohn, Balingen, Germany). The feeding bouts were recorded using a mobile phone camera (Huawei Nexus 6P, 1440 × 2560 px, 120 fps). The recordings were used to determine the time taken for ingestion. Ingestion time was defined as the time from when the bee first touched the sucrose solution with its proboscis until the bee stopped drinking. For each bee, we calculated the absolute ingestion rate i (mg/s):

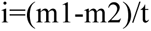

where i is the absolute ingestion rate of a bee, m1 is the mass of the feeder before the foraging bout, m2 is the mass of the feeder after the foraging bout, and t is the ingestion time of the bee. Upon the completion of the test, the bee was sacrificed by freezing and stored in 70% ethanol at -20°C. We measured the intertegular distance (ITD) and the length of the glossa of each bee with a digital calliper (RS PRO Digital Caliper, 0.01 mm ± 0.03 mm) under a dissecting microscope. We then adjusted the absolute ingestion rate i to account for individual size variability using the formula:

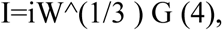

where i is the absolute ingestion rate of a bee, G is the length of the glossa and W is the intertegular distance. This is an adaptation of the formula developed earlier (4) with intertegular distance instead of weight, as it has been shown to be precise at estimating bumblebee weights (5).

To control for evaporation, we located an additional Eppendorf with 50% sugar solution on the opposite side of the test chamber and recorded its weight pre-and post-test for an individual bee. This loss of mass due to evaporation was subtracted from the mass of the test feed after the foraging bout.

### Video analysis

Video analysis was done using BORIS© (Behavior Observation Research Interactive Software, version 7.10.2107 (6). In the judgment bias experiment, we coded two behaviours for each bee. The first behaviour, “Choice”, indicated bee entry into a reward chamber and was classified as a point event, an event which happen at a single point in time. The second coded behaviour, “Latency to choose”, was the time of making the choice and was classified as a state event, i.e., an ongoing event with a duration. For the foraging motivation experiment, we coded a single behaviour, “Drinking duration”, which was classified as a state event that indicated ingestion time.

### Statistical analysis

Our hypothesis and statistical analyses of the main active choice experiment were preregistered at aspredicted.com (#62198). The data were plotted and analyzed using RStudio v.3.2.2 (The R Foundation for Statistical Computing, Vienna, Austria, http://www.r-project.org) and custom-written scripts. To determine the final sample size needed, we used a Bayes Factor approach implemented with the brms package in R (1–3). Prior beliefs about the parameters were specified using a normal distribution with mean 0 and standard deviation 1. Data collection was stopped when the Bayes Factor ≥ 3 (indicating moderate support for HA (2)). All subsequent statistical models for the data were fit by maximum likelihood estimation and, when necessary, optimized with the iterative algorithms BOBYQA. In each analysis, several models were run and compared using the model.sel function in the MuMIn package (38) to select the most appropriate model based on the Akaike information criterion (AIC) scores. We considered the model with the lowest AIC score the best model, i.e., the model that provides a satisfactory explanation of the variation in the data (39). We used the package DHARMa (40) for residual testing of all models.

For the judgment bias analysis, we used the probability of an optimistic choice as the dependent variable, coding choices of reward chambers previously associated with high-value and low -value cues as 1 and 0 respectively. We fit a generalized linear mixed-effect model (GLMM) using the *glmer* function of the *lme4* package with binomial errors and a logit link function (41). The explanatory variables included in the model were “*Treatment*” (categorical: *Control*, *Shaken*, *Trapped*) and “*Cue*” (continuous: 1-5, where 1 = high and 5 = low value cue) which refers to the color displayed on the screen. The identity of the bee (“*ID*”) was included as a random intercept variable.

For the analysis of the choice latency in the judgment bias test, we fit a linear mixed-effect model (LMEM) using the *lmer* function of the *lme4* package (41). To normalize the error distribution, latency data were natural log-transformed and latencies greater than 1.5 times the Inter Quartile Range were excluded (42). The explanatory variables included in the model were “*Treatment*” (categorical: *Control*, *Shaken*, *Trapped*) and “*Cue*” (continuous: 1-5, where 1 = high and 5 = low value cue). In addition, since we expected that optimistic responses would be faster, we also included “*Response Type*” (coded as 1 for optimistic responses, and 0 for pessimistic responses) as an explanatory variable in the model selection process. Bee identity (“*ID*”) was included as a random intercept variable.

In addition to the above models, we ran other statistical tests for some analyses. Data for these tests were first tested for normality and the appropriate tests were subsequently employed for analysis. We ran a one-way ANOVA on the adjusted body size ingestion rate data to test for differences between treatment (Control, Shaking, Trapping). We also used Kruskal-Wallis tests to compare the average number of trials to the criterion in the training phase for different treatment groups, and to investigate the potential impact of the side and color associated with a high-value cue on learning.

### Signal Detection Theory model

We examined whether the behavior of the bees could be modelled with standard signal detection theory, and what could then be inferred about the underlying mechanisms. We assumed that bees learn to make their foraging decisions during training based on the value of an internal signal that is affected by noise. When this signal exceeds an internal decision boundary, the bees behave appropriately for the low reward situation and when it is less than the boundary, they behave appropriately for the high reward situation. We modelled the distribution of the noisy signal and derived the probability of an optimistic response. We fit this model to our data and obtained the decision boundary and the noise for an optimal response given the reward values we used. We compared this decision boundary to the middle value of our response variable. If the boundary was shifted to the right or left of the middle, this would indicate optimistic or pessimistic behavior respectively.

We assumed that bees learn to make their foraging decision during training based on the value of an internal signal x which indicates whether they are in a high or low reward situation. We specified x as a “low reward signal” which has a high value when the cue indicates a low reward. We assumed that bees have some internal decision boundary B, such that when x>B, they behave appropriately for the low-reward situation, and conversely when x<B for the high-reward. Although on average the value of x reflects the cue, it is affected by noise, explaining why bees do not always make the same decision in the same experimental situation.

Since we have fitted our data with a logistic link function, we modelled the distribution of the noisy signal as the first derivative of a logistic function. The standard logistic is

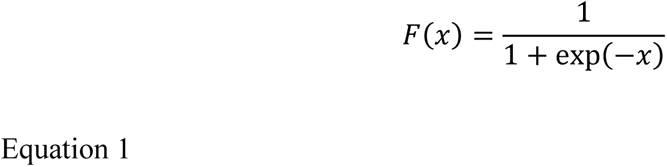

and its first derivative is

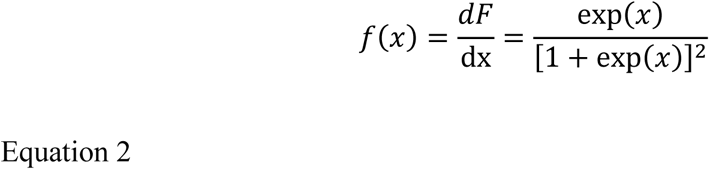

which is therefore the distribution we assume for our noise. This closely resembles a Gaussian distribution with the same standard deviation but has more weight both at the centre and at the tails.

The probability density function governing the distribution of the signal x is 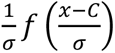, where C represents the value of the cue and s is the noise. The probability of an optimistic response on any given trial is the probability that the value of x on this trial is less than the decision boundary B, given the value of the cue on this trial. This is

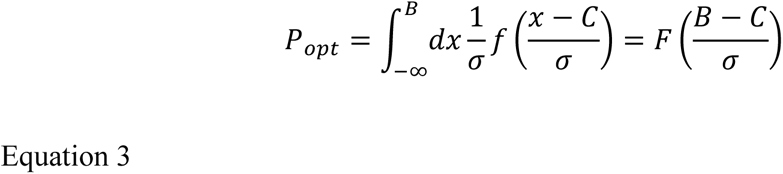

The bee’s behaviour is thus influenced by the noise σ and the decision boundary B. The noise may vary depending on factors like fatigue or attention, while the decision boundary may reflect a cognitive strategy. A common assumption is that the decision boundary is chosen so as to maximise expected reward.

During training, the expected reward is

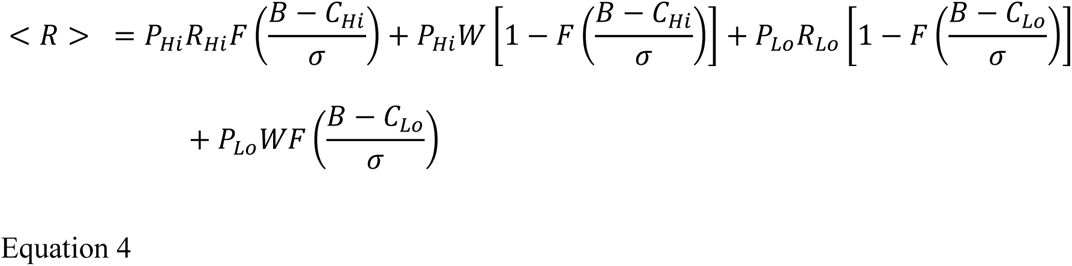

where *P*_Hi_ and *P*_Lo_ represent the probabilities that a given trial offers high or low rewards, *R*_Hi_ and *R*_Lo_ represent the utility to the bee of the 50% and 30% sucrose offered on high or low trials, and W represents the utility of the water obtained when the bee makes the wrong choice.

The optimal boundary *B*_opt_, that maximises the expected reward then satisfies the equation

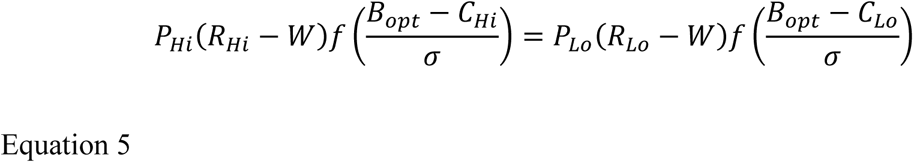

(found by taking the derivative of the expected reward, Equation 4, with respect to B and finding where this is equal to 0). Note that it is possible that the bee isn’t maximising expected reward itself, but some transform of the reward (e.g. reward squared). Since our model has only two values for reward (High and Low), we can still represent any transform as two values (R_Hi_ and R_Lo_) and the model would not be affected by non-linear transforms.

Equation 5 has a simple graphical interpretation. First, the probability distributions for high and low reward are rescaled by their prior probability and by the additional utility of getting the trial right, compared to the water available with the wrong decision. Then, the optimal boundary is where these rescaled distributions cross over (Fig. 4). If the priors and reward utilities were equal, i.e.*P_*H*i_*(*R_*H*i_* − *W*) =*P*_*Lo*_(*R*_*Lo*_ − *W*), then the optimal decision boundary would be exactly in the middle between the two cues values: *B_opt_* = 0.5(*C_*H*i_* + *C*_*Lo*_). If the boundary was shifted to the right or left of the middle, this would indicate optimistic or pessimistic behaviour.

### Drift Diffusion model

Drift diffusion models help shed light on the cognitive processes underlying decision making in choice tasks (43). They help generate estimates of the time taken to accumulate sensory evidence for a particular response and the evidentiary threshold at which the response decision is made. By applying this framework to our experiment, we attempted to see if we could identify which of these two criteria (or both) were changed due to our stress manipulations.

We fit a drift diffusion model to the choice latency data in our three treatments using the R package rtdists (44). The model assumes that the bee accumulates sensory evidence towards a decision and makes the optimistic or pessimistic choice once the evidence has passed a threshold. The thresholds for the pessimistic and optimistic choices were defined to be at 0 and 1 respectively. The decision variable was assumed to begin from a start point *z* somewhere between the two boundaries. It was subject to random noise represented by the diffusion constant *s* but had a drift rate *v* towards one or the other boundary, based on the sensory evidence. In our experiment, *v* should be positive for Cue=1 and negative for Cue=5. In our model, we assumed that v was a linear function of Cue.

## Results

Bumblebees were trained to associate cues of one color with a location containing a high reward of 50% sucrose solution and cues of another color with another location containing lower reward of 30% sucrose solution. The association of rewards with the cue colors and the locations were counterbalanced across all the bees. Bees then experienced one of three treatment conditions. Two groups of bees were physically stressed by shaking or trapping, while the third group served as a control. We then presented the bees with cues of ambiguous colors between the two learnt colors in tests and noted whether they chose the location previously associated with high or lower rewards. We also presented the bee with the cues of the learnt colors during the tests and noted their choices. All the tests were unrewarded and only offered distilled water in the previously rewarding locations.

### Training

During training, a total of 48 bumblebees achieved the learning criterion (80% correct on the last 20 choices) and continued to the judgment bias test. Bees completed training within a minimum of 30 and a maximum of 60 trials. There were no significant differences in the number of trials required to reach the criterion among bees that experienced the high reward on the right or left location (Kruskal-Wallis test: χ^2^ = 2.94, df = 1, p = 0.09). Similarly, there was no difference in the total number of trials to criterion for bees that experienced blue or green as the high reward color (Kruskal-Wallis test: χ^2^ = 0.94, df = 1, p = 0.33). The number of trials required to achieve the learning criterion also did not differ among bees used in each of the three treatment groups (Kruskal-Wallis test: χ^2^ = 0.88, df = 2, p = 0.64).

Bees took significantly longer to choose a low-reward cue in the last choices of the training phase (Table S2, LMEM, Estimate ± standard error = 0.59±0.09, *t* = 6.79, *p* < 0.001). The median latency for choosing in low reward cue trials was 32.2 s (IQR: 35.8), while that for the high reward cue trials was 17.3 s (IQR: 7.34). Thus, bees could differentiate between both the colour cues and the two rewards.

### Physically stressed bees are less optimistic

The best model for our data included the main effects of cue color and treatment (shaking, trapping and control) but not an interaction effect (see supplementary Table S1 for model selection details). Shaking significantly reduced the probability of bees responding optimistically, i.e., choosing the location associated with a high reward (Fig. 2A, Table S2, GLMM, Estimate ± standard error = -1.49 ± 0.57, z = -2.61, *p* < 0.01). Trapping with a robotic arm also significantly reduced the likelihood of an optimistic response (Fig. 2A, Table S2, GLMM, Estimate ± standard error = -1.26 ± 0.56, z = -2.23, *p* = 0.026). Bees were also significantly less likely to respond optimistically to cues with colors further away from that of the high reward cue (Fig. 2A, Table S2, GLMM, Estimate ± standard error = -1.79 ± 0.21, z = -8.39, *p* < 0.001).

**Figure 2.**
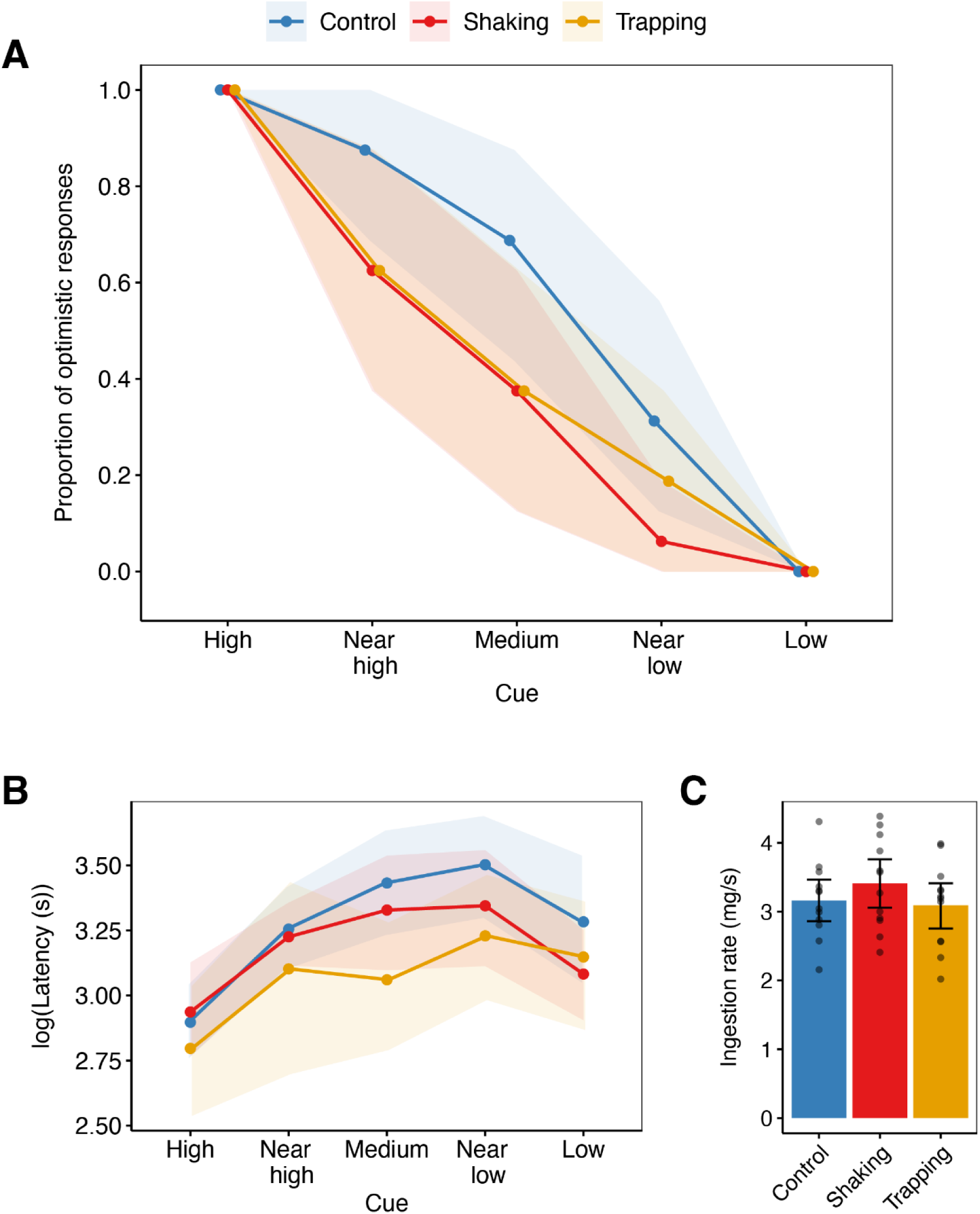
Bee responses to test cues. **A)** Proportion of bees (N = 16 per treatment) making an optimistic choice (choosing a reward chamber associated with a high reward) in response to each of five cues. **B)** Latency of making the choice in response to each of five cue values (N = 16 bees per treatment). **C)** Average ingestion rate of high reward (50% sugar solution) for bees in each treatment group (N = 12 bees per treatment). The treatment groups were control (blue), shaking (red), and trapping (orange). The test cues were high, near high, medium, near low, and low value cues depending on their distance to the colors of high-and low-reward cues. Points and bars represent means, and the shaded areas and error bars represent 95% bootstrapped confidence intervals. Dots represent values form individual bees.

### Feeding motivation and choice latencies

We tested the ingestion rate of sucrose solution as a measure of the feeding motivation of the bees. The mean (± s.d.) ingestion rate by shaken and trapped bees was 3.42 ± 0.67 mg/s, and 3.17 ± 0.61 mg/s respectively. The mean ingestion rate observed in control bees was 3.17 ± 0.55 mg/s. These rates did not differ significantly between treatment groups (Fig. 2C, ANOVA: F(2, 33) = 0.642, p = 0.533).

We also examined the change in the latency to make a choice in the experiments. The best-fitting model included treatment, cue value and response type (optimistic or pessimistic) as fixed predictors and an interaction between cue value and response type (supplementary Table S1). Bees in the Trapping treatment were significantly faster to make a choice than control bees (Fig. 2B, Table S2, LMEM, Estimate ± standard error = -0.23 ± 0.1, t value = -2.25, p = 0.029) but were not faster than those in the Shaking treatment (Fig. 2B, Table S2, LMEM, Estimate ± standard error = - 0.12 ± 0.1, t value = -1.15, p = 0.256). Shaken bees were not significantly faster to make their choices than control bees (Fig. 2B, Table S2, LMEM, Estimate ± standard error = -0.11 ± 0.10, t value = -1.121, p = 0.27). All bees were also significantly slower to make a choice when the cue color was further away from that of the high reward cue (LMEM, Estimate ± standard error = -0.09 ± 0.03, t value = - 2.6, p < 0.01). Finally, bees were faster when making optimistic choices compared to pessimistic ones (LMEM, Model Estimate ± standard error = -0.93 ± 0.16, t = -5.74, p < 0.001).

### Signal-detection theory model

According to a standard signal-detection theoretic approach, the probability that a bee makes an optimistic choice for Cue level *C* is (Equation 3)

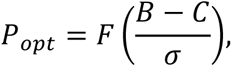

where **σ** is the noise on the internal signal, *B* is the decision boundary, and F is the logistic function. This is exactly the model fitted by our generalized linear mixed model (GLMM, see above), with the fitted gradient for *Cue* corresponding to −1/*σ* and the intercept corresponding to *B*/*σ*. Thus, the fact that we found no interaction between *Cue* and *Treatment* indicates that the effective noise level is not changed by our manipulations. The estimate of -1.79 for the gradient (Table S2) allows us to infer an effective noise level of **σ** = 0.56, in our units where Cue runs from 1 (high reward) to 5 (low reward).

However, the significant main effect of *Treatment* indicates that the decision boundary was different in the two cases. The estimate of 6.05 (Table S2) for the intercept in the control condition implies that the decision boundary in this condition is 3.38. Bees in the Control treatment (Fig. 2A) are thus equally likely to make the optimistic or pessimistic response when the cue is a little closer to “near low” than medium (3). The fact that the intercept drops by -1.49 for the Shaking treatment and -1.26 for Trapping (Table S2) implies that the boundary shifts leftward to 2.55 and 2.68, respectively, in these conditions. The point at which these bees are equally likely to make optimistic and pessimistic choices is closer to “near high” than to medium (Fig. 3B).

**Figure 3.**
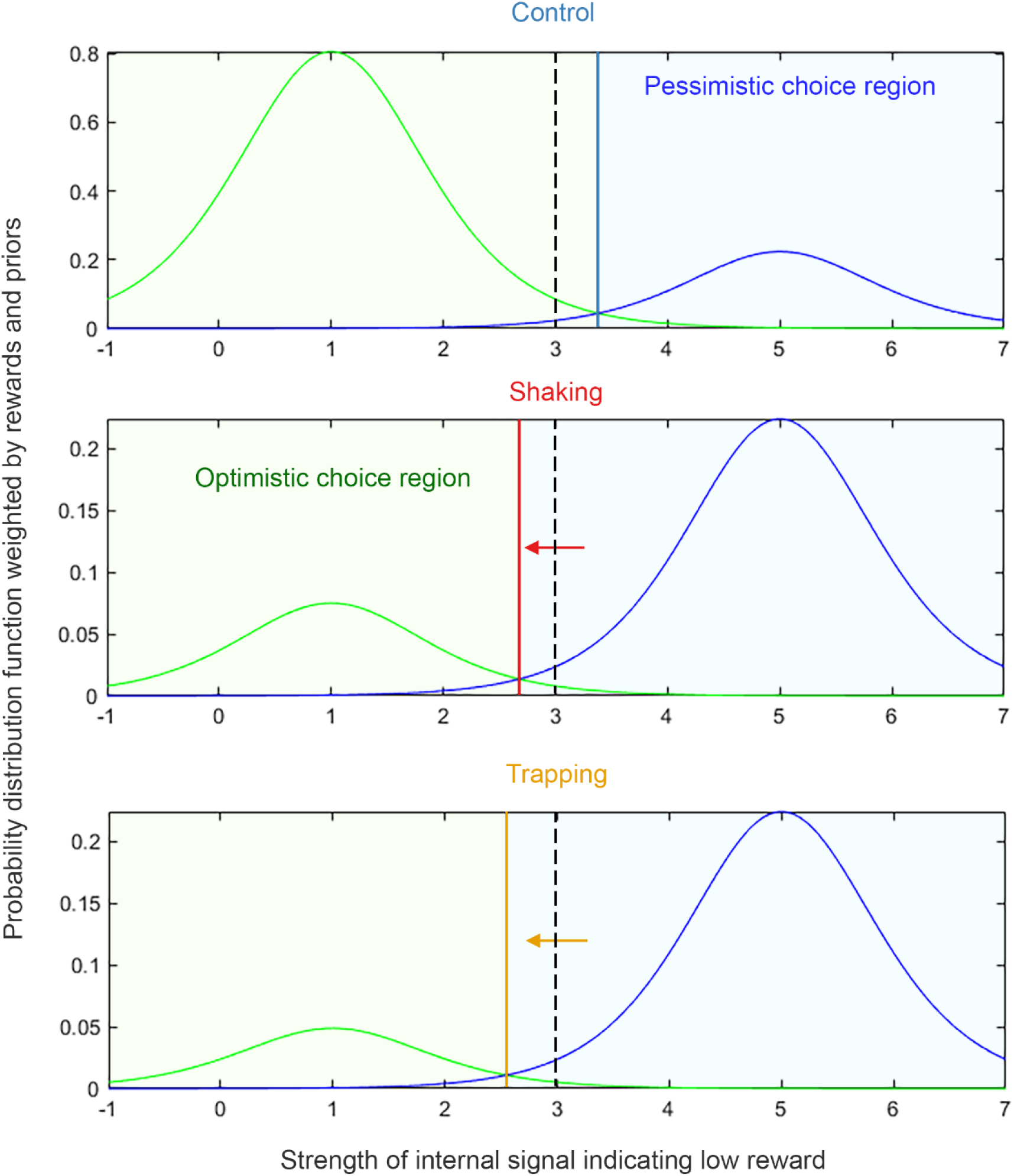
Bee decision-making boundaries and priors fitted by a signal-detection model. Curves depict the probability density functions for responses based on the internal signal *x* indicating a low reward. In each case, the original distribution has been weighted by the product of the value of that reward and its probability of occurring (see methods, Equation 5). The two curves in each panel depict the probabilities that the cue indicates high reward (green, centred on 1) or low reward (blue, centred on 5). Solid lines depict the decision boundary B inferred from the Generalized Linear Mixed Model fit to our data. Dotted lines indicate the medium point for comparison. Regions to the right of the solid boundary line are regions where the bee makes pessimistic choices (shaded blue). Regions to the left are regions where the bee makes optimistic choices (shaded green). Arrows depict the shift in boundaries compared to the control condition. The three panels depict the conditions for the Control (top), Shaking (middle) and Trapping (bottom) treatments. Note the change in axes in the lower two panels.

**Figure 4.**
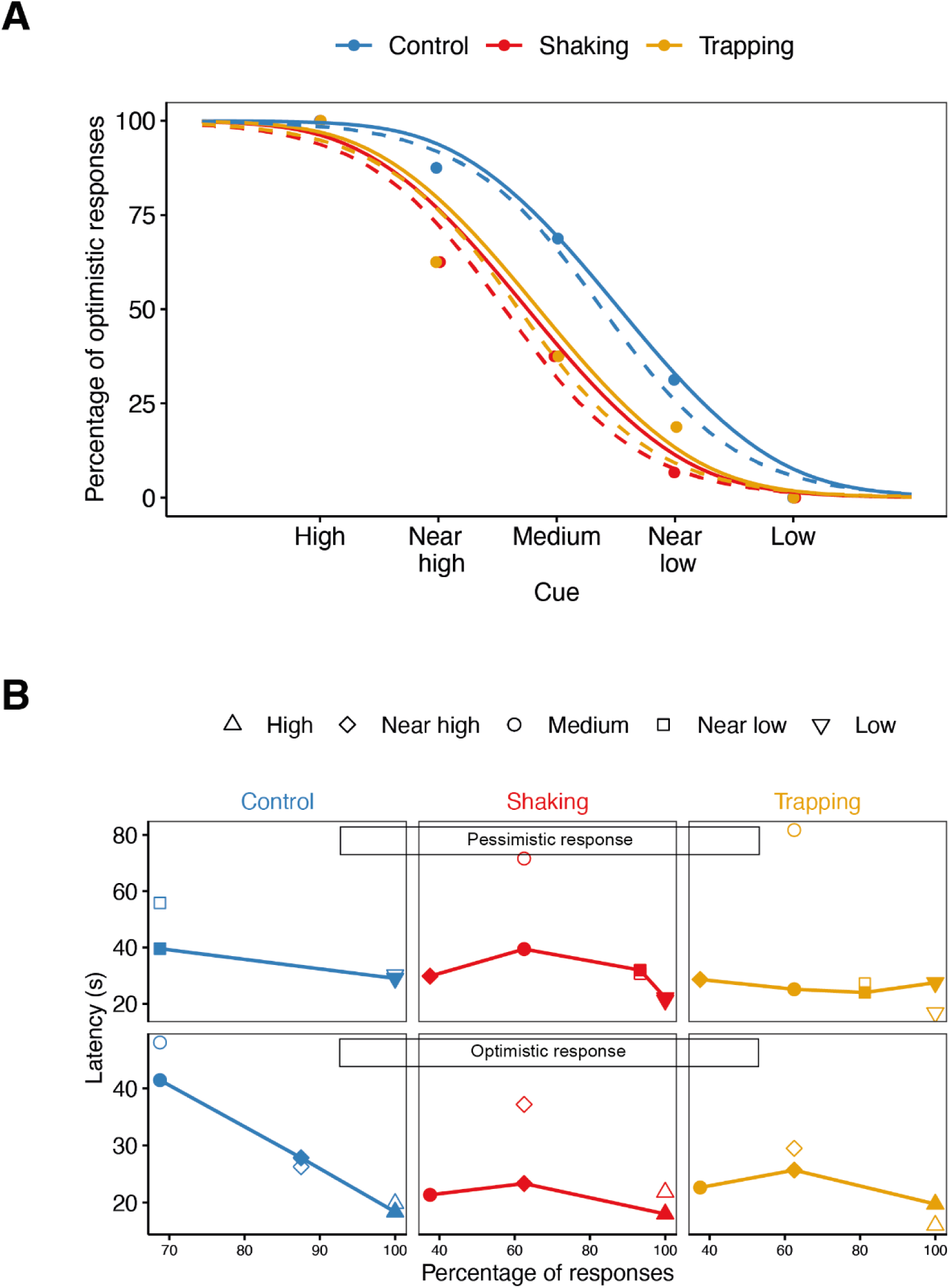
Drift diffusion model. **A)** Proportion of optimistic choices made by the bees in each treatment in response to the different cues. Points show the data, dashed curves show the predictions of a fitted logistic regression model with main effects of Treatment and Cue but no interaction. Solid curves show predictions of a fitted drift diffusion model. Colours depict the different treatments: Control (blue lines), Shaking (red lines) and Trapping (orange lines). **B)** Drift diffusion model fit to latencies. Filled symbols linked with lines show median latencies as a function of the percentage of responses made, for pessimistic (top) and optimistic (bottom) responses in the three treatments (columns). Empty symbols show predictions of the fitted drift diffusion model. Symbols show Cue value. There is a high percentage of optimistic responses for high (triangles) and near high (diamonds) cues and a high proportion of pessimistic responses for low (inverted triangles) and near low (squares) cues.

In our fitted model, weighted probability distributions for both low and high rewards have an equal spread, reflecting the noise level inferred from the GLMM. In the Control treatment, the shift of the decision boundary reflects the greater weight given to the high reward. Quantitatively, the extent of the shift, together with the fitted noise level, implies that the high reward is given 3.6 times the weight of the low reward. This result also cannot be explained merely by the bees not perceiving the medium colour as midway between blue and green since both the high and low reward trials combine data from trials where the cue was blue and trials where it was green. Instead, this result might, for example, indicate that the bees understand that both rewards are equally likely (P_Hi_ = 50%) and find the 50% sucrose solution 3.6 times as rewarding, relative to water, as the 30% solution.

The fact that the decision boundary is to the left of neutral in the Shaking and Trapping treatments indicates that here, greater weight is given to the low reward (Fig. 3B). Assuming we can discount the possibility that the reward value has inverted (i.e., that stressed bees find 30% sucrose more rewarding than 50%), this must represent a shift in the priors, such that stressed bees now consider high-reward trials less likely. To match the extent of the leftward shift, given the noise level inferred from our GLMM fit, the low reward must be weighted 4.6 times as much as the high reward. If the reward ratio were 3.6, this would imply that the bees behave as if the perceived probability of the high reward was 6%. However, if stressed bees find 50% and 30% sucrose equally valuable, i.e., the stress has removed the difference in reward utility, then the observed shift in decision boundary could be produced with a less dramatic shift in the priors, with perceived probability of the high reward being 18%.

### Drift diffusion model

Drift diffusion models generate estimates of the time taken to accumulate sensory evidence for a particular response and the evidentiary threshold at which the response decision is made. By applying this framework to our experiment, we attempted to see if we could identify which of these two criteria (or both) were changed due to our stress manipulations. Our best model (as indicated by the Akaike Information Criterion) was obtained by allowing the time prior to making a decision and the value of the drift rate for Cue = 3 (v3) to vary between treatments, while fitting all data with the same values for the diffusion constant s, start point zr, the dependence of drift rate on cue, vGradient, and noise on the drift rate, sv. The drift diffusion model predicts not only the bees’ choices (Fig. 4A) but also the latencies for both optimistic and pessimistic choices (Fig. 4B). There are not enough trials to accurately estimate the latency distributions (just 16 trials for each Cue/Treatment combination, thus < 16 for each choice). The model for latencies is, therefore, not a good fit (Fig. 4B), and it would be unwise to draw too strong conclusions from this fitting effort. Nevertheless, the fitted model implies a few key points.

Firstly, by default, bees tend to be biased towards the more rewarding choice. The start point for the decision variable is not midway between the two boundaries, 0.5, but closer to the boundary for the optimistic choice, 0.56. As noted in the signal detection theory model, being biased towards the high-reward condition helps to maximise the expected reward. Secondly, stress did not affect sensory noise. As in the logistic regression model, we found that the best model was obtained by assuming that sensory noise, here represented by the diffusion constant s, was the same for all groups. Thirdly, stressed bees spend less time on non-decision activity: the model fitted more time on non-decision activity (e.g., flying across the arena) for the control bees than for the shaken or trapped bees. This could perhaps suggest that stressed bees might not want to spend time exploring what could potentially be a dangerous environment. Finally, this model also confirms that the stressed bees are more pessimistic. This is shown by the fitted drift rate for the medium cue, Cue = 3. In the absence of bias, the drift rate should have been zero in this case, since the cue was designed to be exactly midway between the high and low reward cues (and counterbalancing ensured that it was on average). Control bees nevertheless showed a small positive drift rate for this cue, indicating that they took it as weak evidence for a high reward. As noted above, this bias towards high reward helps maximise expected reward. However, shaken and trapped bees both showed a small negative drift rate, indicating perceived weak evidence for low reward. This is what accounts for the leftward shift in the response curves for stressed bees. Note that even though, according to the model, all bees start slightly biased towards a high-reward response (z = 0.55), in stressed bees, the negative drift rate for the medium cue is enough to bias responses towards the pessimistic response.

## Discussion

We developed a novel task to assess emotion-like states in bees. Using an active choice judgment bias task, we demonstrated that physically stressed bees are more likely to make pessimistic choices when faced with ambiguous stimuli. A signal detection model of our data suggests that this behaviour is explained by a reduced expectation of rewards. We thus provide strong evidence for bee judgement biases and a possible explanation for bee behaviour in judgement bias tasks.

Most studies of judgement bias tests have used a go/no-go paradigm. The results of these studies can be challenging to interpret due to confounds from other factors that do not involve stimulus judgements such as, for example, motivation. Our active choice design avoids these complications. Motivation alone cannot therefore explain the observed shift in responses in the manipulated bees in our experiment. This is further supported by the results of our ingestion rate experiment, where we do not find differences in feeding motivation. Only one previous study has used an active choice design to study judgement biases in insects (5). In that study, flies had to choose between two odours, one associated with a reward and another with punishment. Rather than using reward and punishment, we developed a novel paradigm for insects that uses two rewards of different quality. This allowed us to investigate the mechanisms underlying the judgement bias in further detail and test how negative states modulate expectations and perceptions of reward. Using previous paradigms involving reward and punishment as the expected outcome can make it easier to detect affect-dependent judgement bias (23). We, however, find a bias in bee behaviour when using two rewards and an active choice paradigm, providing stronger evidence for affect-dependent processing in insects.

### Bees learnt the stimulus-outcome associations

When performing an active choice task, it is important to ensure that the rewards used to condition the animals’ responses are not perceived as equally favourable. If so, the results of tests using ambiguous stimuli would reflect the animal’s colour preferences rather than its interpretation of the outcome associated with a particular colour. Bumblebees, however, can use colour cues to discriminate between rewards of varying value and prefer higher concentrations of sugar solution, including the colours and concentrations we used in our experiments (24). In our experiments, too, the bees chose high rewards significantly faster than lower rewards at the end of the training phase. In the tests, bees in all treatment groups also made slower choices as the cue value moved further away from the one indicating a high reward. The shorter choice latency towards the high reward cue suggests that bees maintain their preference for higher rewards even after experiencing stress. This demonstrates that the bees distinguished between the high and low rewards, regardless of the associated colour.

### Physical stress was not detrimental to bee sensory perception

Manipulations in judgement bias tasks need to change decision-making without impairing sensory perception or discrimination. In one previous test of judgement biases, shaken honeybees showed a decreased response not only to ambiguous odour mixtures but also to the conditioned negative odour (4). This decrease has been suggested to indicate an improved ability to differentiate odours rather than a negative bias in judgement (10). In our experiment, however, the bees were perfectly accurate when responding to both conditioned cues (high and low) in the tests. The drift diffusion model further indicates that the stress treatments did not change the sensory noise. Our manipulations thus did not impair the colour discrimination abilities and memory of the bees. The preservation of high colour discrimination abilities is not surprising, as previous studies on Drosophila have successfully used shaking in aversive learning paradigms (25). Similar trapping mechanisms to the ones we used have also been employed in aversive learning tasks in bees (26).

### Active choices are better indicators of judgments than latencies

Latency is often used in go/no-go judgment bias tests to evaluate the emotional states of animals (6). When evaluating an emotional state, it is important to determine whether it is positive or negative (known as valence). However, relying solely on latency as a measure of valence is not always reliable, as it can be affected by other factors unrelated to emotions. An increase in approach latency may be associated with a general increase in reactivity and arousal, for example, due to the increased energetic demands after experiencing stressful events (27). It may also indicate a shift in the perceived value of the reward and differences in motivation (28). Relying solely on latency can therefore make it challenging to interpret the results of judgment bias tests. For instance, exposure to a positive event has been reported to cause both longer (29) and shorter (30) response times to ambiguous stimuli.

Only one study has used latencies to measure emotion-like states in bees (6). This study used a go/no-go type of judgment bias test to demonstrate an optimistic bias in bumblebees after receiving an unexpected reward of sugar solution. As predicted, unexpected rewards reduced the latency with which bees approached ambiguous stimuli. However, the treatment also caused an increase in thoracic temperature which has been linked to increased motivation for foraging in other studies (31). Further experiments did indicate that optimism was a more plausible explanation, but choice latency clearly could be influenced by motivational changes as well as judgements. Our results showed that after trapping, bees had shorter latencies than the control bees. This could, in principle, have indicated a positive state, again demonstrating the difficulty of using latencies alone to interpret judgement bias data. However, since our study was an active choice design, we could more reliably use the choices made by the bees rather than their latencies. Choices can better indicate affective valence, showing that the trapped bees were in a pessimistic state in our study. This makes a strong argument in favour of active choice judgement bias tasks such as the one we used in our study.

### Pessimistic choices by bees are related to a significant change in prior expectations

To unravel the potential mechanisms underlying the choices made by the bees, we employed a signal detection approach, which has been previously suggested as a valuable tool for investigating affective biases (32). A recent study has suggested that judgement biases in bees may be caused by a shift in stimulus-response curves (7). However, this study did not investigate the underlying causal mechanisms of this shift. In our model, the estimation of future outcomes combines estimates of the probability of an outcome occurring and the magnitude of the payoff from an outcome. Both the signal detection and drift diffusion analyses demonstrate that control bees exhibit a higher probability of responding optimistically to ambiguous cues, indicating an expectation of high rewards. Such a bias would not be suboptimal as it is in fact what is predicted by a rational, fully informed strategy which optimises expected reward. Even if the bees are estimating the priors correctly as 50-50, the difference in reward utility will still shift the decision boundary towards the cue indicating low reward (Fig. 4A). Our model shows that the control bees are behaving as if 50% sucrose is 3.6 times more valuable, relative to water than 30% sucrose. Thus, the data admit the possibility that the bees’ behaviour is completely rational and unbiased, and the 50% sucrose is much more rewarding.

However, the decision boundary and drift rate for the stressed bees are harder to interpret. Here, the decision boundary is to the *left* of neutral and the drift rate is negative. Previous studies have shown that acute stress can increase an animal’s sensitivity to the reward (33). However, the observed left shift of the decision boundary in stressed bees cannot plausibly reflect such a change in reward sensitivity since a leftward shift could only be produced if the value of high and low rewards were swapped, i.e., if 50% sucrose became less rewarding than 30%. However, it could reflect a pessimistic bias in expectations, i.e., that the stressed bees behave as if high-reward priors are less likely (*P*_Hi_ < *P*_Lo_), perhaps because in nature high rewards are indeed scarcer when conditions are stressful. This can account for a leftward shift, but the large quantitative extent of the shift is still surprising. Since the noise remains relatively small, as indicated by the perfect performance for high and low cues, we have to postulate enormous changes in the priors to produce the observed shift. To obtain the decision boundary of 2.55 inferred for shaken bees, we would have to postulate that shaken bees estimate P_Lo_ = 94%, i.e., they expect a high reward to be available on only one trial in 20. This assumes that the reward utility remains the same, with a high reward 3.6 times as valuable as a low. If the relative utility of the high reward increased, e.g., because of an increased need for sucrose after stress (27), the priors would have to shift even further from 50%. However, one possibility is that, contrary to the assumptions of our model, the noise was not uniform for all cues, and there was more sensory noise on intermediate values of the cue. If this were so, the change in priors would not need to be as dramatic, although the basic result of changed priors would remain true.

By employing an active choice judgment bias task, our results further support the possibility of emotion-like states in bees and suggest that these states could be found across very different animals. By implementing a more demanding active choice design, we provide robust evidence that neither motivational factors nor colour discrimination alone can account for the observed cognitive biases. Importantly, our modelling indicates that the pessimistic-like behaviour displayed by bees in a negative state represents a significant shift in their prior expectations of rewards. These insights offer the first analytical models of the underlying causal mechanisms of state-dependent judgment biases in insects, opening up new avenues for exploring state-dependent decision-making in insects.

## Acknowledgments

This work is supported by a BBSRC David Phillips fellowship BB/S009760/1 to VN. OP is supported by a PhD scholarship from the Faculty of Medical Sciences, Newcastle University. We are grateful to Théo Robert for insightful discussions.

## Data Availability Statement

All relevant data and code used for analysis to support this paper are available as supporting information.

