## Supplemental Tables for "Physically stressed bees expect less reward in an active choice judgement bias test"

**Supplementary Information**

Table S1. Summary of model selection to analyse the impact of stress treatments (shaking/trapping) on performance in the judgement bias task

| | Explanatory variables | | | |
|---|---|---|---|---|
| Dependent variable | Fixed | Random | d.f. | AIC |
| Optimistic Response | **Treatment+Cue** | **ID** | **5** | **179.81** |
| | Treatment*Cue | ID | 7 | 183.24 |
| | Cue | ID | 3 | 184.39 |
| | Treatment | ID | 4 | 335.19 |
| log(Choice latency) | **Cue*Response+Treatment** | **ID** | **8** | **234.64** |
| | Cue*Response | ID | 6 | 235.46 |
| | Cue+Treatment+Response | ID | 14 | 242.35 |
| | Cue*Treatment*Response | ID | 6 | 256.38 |
| | Response | ID | 4 | 256.47 |
| | Cue+Response | ID | 5 | 257.39 |
| | Cue+Treatment+Response | ID | 7 | 257.46 |
| | Cue*Treatment+Response | ID | 9 | 259.14 |
| | Cue+Treatment*Response | ID | 8 | 260.15 |
| | Cue | ID | 4 | 260.17 |
| | Cue+Treatment | ID | 6 | 260.92 |
| | Cue+Treatment*Response | ID | 9 | 261.28 |
| | Cue*Treatment | ID | 8 | 262.74 |
| | (1\|ID) | ID | 3 | 273.35 |
| log(Training latency) | **Cue** | **ID** | **4** | **137.36** |
| | Cue+Treatment | ID | 6 | 144.68 |
| | Cue*Treatment | ID | 8 | 150.98 |
| | (1\|ID) | ID | 3 | 162.20 |
| | Treatment | ID | 5 | 169.53 |

Three models were fit to analyse 1) the likelihood of choosing reward chamber associated with high reward (optimistic response) and 2) Choice latency in the test and 3) Choice latency during training in trials with high and low cues. Only summaries of the best fit models are shown. For each model, fixed and random explanatory variables, degrees of freedom (d.f.) and Akaike's Information criterion (AIC) are detailed. For each dependent variable, the selected model, i.e., the one with the lowest AIC, is indicated in bold.

Table S2. Summary of the best fit statistical models analysing the impact of stress treatments (shaking/trapping) on performance in the judgement bias task.

| Response variable | | Estimate | Std. Error | z value/ t value | Pr(>\|z\|) |
|---|---|---|---|---|---|
| Optimistic response | (Intercept) | 6.05 | 0.81 | 7.46 | 0.000 |
| | **Cue** | **-1.79** | **0.21** | **-8.39** | **0.000** |
| | **Treatment (Shaking)** | **-1.49** | **0.57** | **-2.61** | **0.009** |
| | **Treatment (Trapping)** | **-1.26** | **0.56** | **-2.23** | **0.026** |
| log(Choice latency) | (Intercept) | 3.77 | 0.16 | 23.42 | 0.000 |
| | **Cue** | **-0.09** | **0.03** | **-2.60** | **0.010** |
| | **Response (Optimistic choice)** | **-0.93** | **0.16** | **-5.74** | **0.000** |
| | Shaking | -0.11 | 0.10 | -1.12 | 0.268 |
| | Trapping | -0.23 | 0.10 | -2.25 | 0.029 |
| | **Cue*Response (Optimistic choice)** | **0.26** | **0.05** | **5.15** | **0.000** |
| log(Training latency) | (Intercept) | 2.87 | 0.08 | 35.95 | 0.000 |
| | **Cue** | **0.59** | **0.09** | **6.79** | **0.000** |

The table provides summaries of the best fit models for the effects of the treatments, shaking and trapping, on the performance of subjects in the judgment bias task. The data is presented in terms of estimated coefficients, standard errors, z-values (Optimistic response model) or t-values (Choice latency model), and the associated p-values for response variables and predictor variables.
